# Discovery of functional alternatively spliced *PKM* transcripts in human cancers

**DOI:** 10.1101/613364

**Authors:** Xiangyu Li, Cheng Zhang, Woonghee Kim, Muhammad Arif, Chunxia Gao, Andreas Hober, David Kotol, Linnéa Strandberg, Björn Forsström, Åsa Sivertsson, Per Oksvold, Morten Grøtli, Yusuke Sato, Haruki Kume, Seishi Ogawa, Jan Boren, Jens Nielsen, Mathias Uhlen, Adil Mardinoglu

**Affiliations:** Science for Life Laboratory, KTH - Royal Institute of Technology, Stockholm, Sweden; Department of Chemistry and Molecular Biology, University of Gothenburg, SE-412 96 Gothenburg, Sweden; Department of Pathology and Tumor Biology, Institute for the Advanced Study of Human Biology (WPI-ASHBi), Kyoto University, Kyoto, Japan; Department of Urology, Graduate School of Medicine, The University of Tokyo, Tokyo, Japan; Department of Medicine, Centre for Hematology and Regenerative Medicine, Karolinska Institute, Stockholm, Sweden; Department of Molecular and Clinical Medicine, University of Gothenburg, Sahlgrenska University Hospital, Gothenburg, Sweden; Department of Biology and Biological Engineering, Chalmers University of Technology, Gothenburg, Sweden; Centre for Host-Microbiome Interactions, Faculty of Dentistry, Oral & Craniofacial Sciences, King’s College London, London, SE1 9RT, United Kingdom

**Keywords:** *PKM*, alternative splicing, transcriptomics, cancer

## Abstract

The association of pyruvate kinase muscle type (*PKM*) with survival of cancer patients is controversial. Here, we focus on different transcripts of *PKM* and investigate the association between their mRNA expression and the clinical survival of the patients in 25 different cancers. We find that the transcript encoding PKM2, and three other functional transcripts are prognostic in multiple cancers. Our integrative analysis shows that the functions of these four transcripts are highly conservative in different cancers. Next, we validate the prognostic effect of these transcripts in an independent kidney renal clear-cell carcinoma (KIRC) cohort and identify a prognostic signature which could distinguish high- and low-risk KIRC patients. Finally, we reveal the functional role of alternatively spliced *PKM* transcripts in KIRC, and discover the protein products of different transcripts of *PKM*. Our analysis demonstrated that alternatively spliced transcripts of not only *PKM* but also other genes should be considered in cancer studies, since it may enable the discovery and targeting of the right protein product for development of the efficient treatment strategies.

## Introduction

Pyruvate kinase muscle type (*PKM*) is the most-studied isoform of pyruvate kinase and catalyzes the final step in glycolysis^1^. It is one of the key mediators of the Warburg effect and plays a pivotal role in controlling tumor metabolism. It has been reported that the mRNA and protein expression of *PKM* is strongly associated with the survival of cancer patients, but the direction of the correlation was contradictory since both activation and inhibition of this enzyme have been suggested for effective treatment of the cancer patients^2^. In the Human Pathology Atlas^3^, high expression of *PKM* is significantly (log-rank *p*-value<0.05) associated with the unfavorable prognoses in liver hepatocellular carcinoma (LIHC), pancreatic adenocarcinoma (PAAD), head and neck squamous cell carcinoma (HNSC) and lung adenocarcinoma (LUAD) whereas it is also associated with favorable prognoses in kidney renal clear-cell carcinoma (KIRC), skin cutaneous melanoma (SKCM), stomach adenocarcinoma (STAD) and thyroid carcinoma (THCA). Thus, mRNA expression of *PKM* has ambiguous indication of patients’ survival in different cancer-types.

The oncological roles of differentially spliced transcripts of *PKM* including PKM1 and PKM2, which are mutually exclusive exons 9 and 10^4^, have been previously investigated. It has been reported that overexpression of PKM1/2 isoforms promotes tumorigenesis or induces poor prognoses of patients in multiple cancers^5-17^ whereas PKM1 expression in place of PKM2 inhibits tumor cell proliferation^18,19^. Moreover, it has been reported that methylation or deletion of PKM2 promotes tumor progression in liver cancer, breast cancer and medulloblastoma^20-23^. Therefore, the function of alternative splicing products of *PKM* in tumor oncogenesis and progression remains controversial.

Due to alternative splicing, there are 14 known isoforms of the *PKM*, of which PKM1 and PKM2 are two major isoforms. To our knowledge, the roles of other protein products of *PKM* apart from PKM1 and PKM2 have not been studied. In this study, we focused on 14 different transcripts of *PKM* and systematically investigated the biological functions of each transcript as well as their association with the clinical outcomes in 25 different cancer types.

## Results

### The prognostic effect of *PKM* at the transcript level

We retrieved mRNA expression of *PKM* and its 14 transcripts together with the clinical survival metadata of patients in 25 different cancer-types in The Cancer Genome Atlas (TCGA) and investigated the associations between the mRNA expression of the transcripts with patients’ survival outcomes (Table S1). We performed a Kaplan-Meier survival analysis for the patients by classifying the patients into two groups with high and low expression of the investigated transcript by optimally selecting a cutoff from the 10^th^ to 90^th^ expression percentiles yielding the lowest log-rank *p*-value as in our previous study^3^ (Table S2). As shown in Figure 1A, the mRNA expression of the *PKM* indicated opposite survival outcomes in different cancer-types. At the transcript level, we found that seven transcripts had mRNA expression (average TPM) > 5 and six of these transcripts, including ENST00000335181 (encoding PKM2), ENST00000319622 (encoding PKM1), ENST00000561609, ENST00000389093, ENST00000568883 and ENST00000562997, are significantly associated with patients’ survival outcome in at least one cancer. Among them, the mRNA expression of transcript encoding PKM2 exhibited a very similar prognostic indication to *PKM* in all cancer types since it represents ∼95% of its mRNA expression (Table S1). Notably, we observed that the expression levels of transcript encoding PKM1, which has been associated with different cancer types^5,6,24,25^ is only prognostic in HNSC.

**Figure 1.**
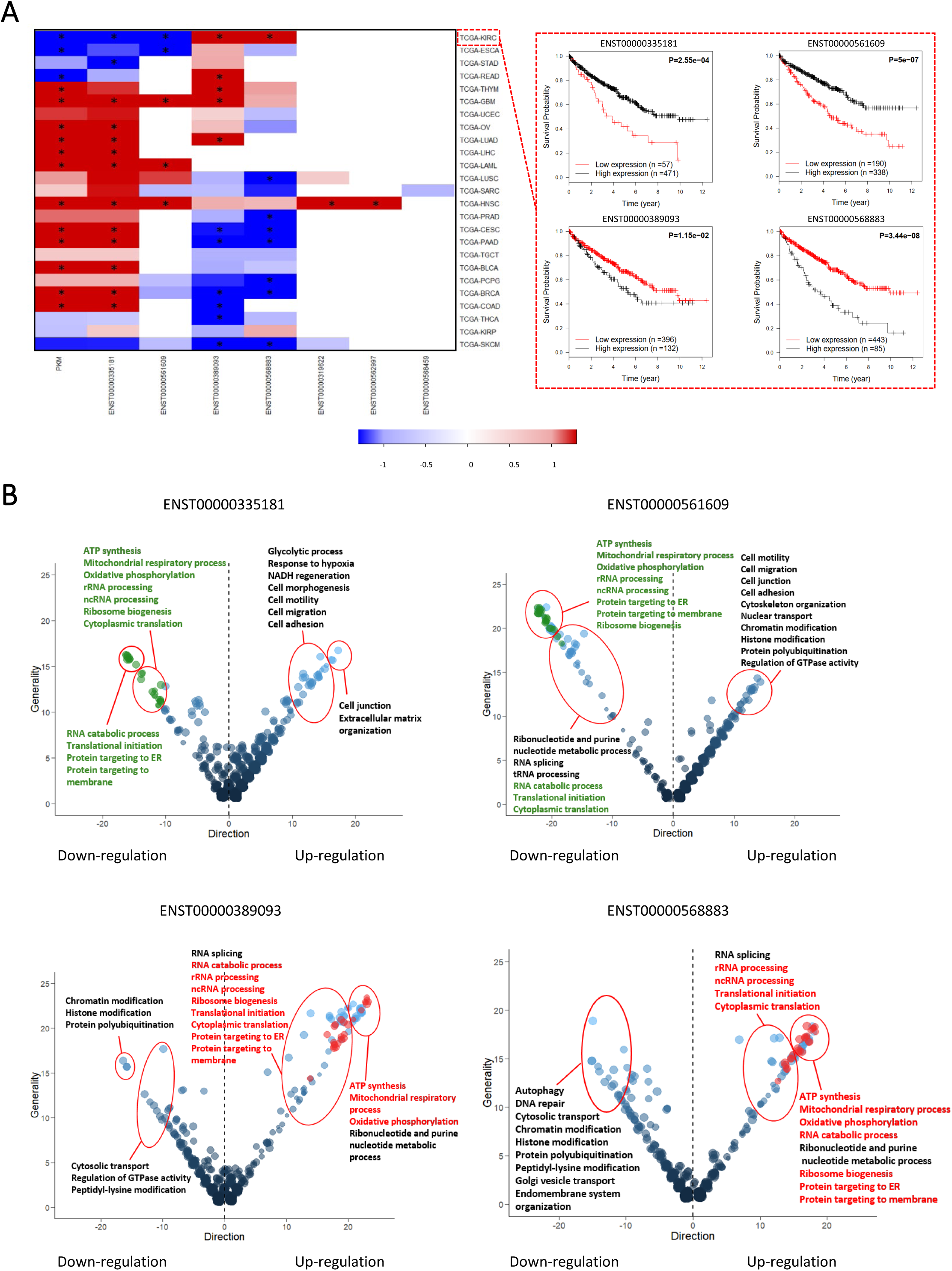
Prognostic and functional analysis of *PKM* and its alternatively spliced transcripts in TCGA data. (A) Heat map of the log-rank *p* values (on the negative log 10 scale) of *PKM* and seven transcripts (average TPM>5) in 25 cancer-types. Six of these transcripts are significantly associated with patients’ survival outcome in at least one cancer. The Kaplan Meier plots for KIRC was exemplified. (B) Bubble plot showing the common enriched Gene Ontology (GO) terms among the 25 cancer-types in the TCGA. Bubble sizes represent numbers of genes associated to the biological function in a specific GO term; the x and y axes indicate the directions and generalities of the GO terms. Generality is defined by the number of cancers with differentially expressed genes (DEGs) associated with each transcript; direction is defined by the number of cancers with their upregulated genes over-representing the GO function minus the number of cancers with downregulated genes over-representing the GO function. Note that only functions with more than ten generalities are shown. The red bubbles denote the commonly detected Go terms enriched with up-regulated DEGs and the green bubbles denote the commonly detected Go terms enriched with down-regulated DEGs. Significantly enriched GO terms for each transcript are provided in Table S3.

High expression of ENST00000335181, ENST00000561609, ENST00000389093 and ENST00000568883 indicated opposite clinical survival outcome in different cancer-types. This is exemplified by ENST00000568883, whose high expression indicates unfavorable survival in KIRC patients and favorable survival in lung squamous cell carcinoma (LUSC), prostate adenocarcinoma (PRAD), cervical squamous cell carcinoma and endocervical adenocarcinoma (CESC), PAAD, pheochromocytoma and paraganglioma (PCPG), breast invasive carcinoma (BRCA) and SKCM. We also found that the expression of *PKM* indicated an opposite survival outcome of patients compared to that of its transcripts in other cancer-types. For example, ENST00000568883 exhibited the opposite prognostic indication compared to *PKM* in KIRC, CESC, PAAD and BRCA (Figure 1A). Moreover, the high expression of different *PKM* transcripts may induce opposite prognoses in patients with the same cancer. This is exemplified in KIRC, where high expression of ENST00000335181 and ENST00000561609 pair indicated favorable prognoses of patients and high expression of ENST00000389093 and ENST00000568883 pair indicated the opposite, and similar scenarios could be found in CESC, PAAD, BRCA and colon carcinoma (COAD).

### Biological functions of the *PKM* transcripts

We identified four different transcripts of *PKM* including ENST00000335181, ENST00000561609,ENST00000389093andENST00000568883which exhibited opposite prognostic effect in multiple cancer types. Hence, we investigated whether this opposite trend is also observed at the functional level in all cancer types. To systematically identify the functions of the four prognostic *PKM* transcripts, we identified the differentially expressed genes (DEGs) between patients with the top 25% high expression and bottom 25% low expression of each transcript in all cancers (FDR < 1.0e-05). We performed a gene ontology (GO) enrichment analysis using the DEGs driven by each transcript and summarized the results of the enriched GO terms for all transcripts in all cancer types (FDR < 0.001, Figure 1B and Table S3). As shown in Figure 1B, if a GO term is enriched with DEGs in multiple cancers, the directionality of the DEGs often follows the same direction. For example, as shown in Figure 1B, the DEGs identified by comparing high and low expression of ENST00000335181 is enriched in extracellular matrix organization pathway in 16 cancers and are always associated with the upregulated genes. Our analysis indicated that although the prognostic effect of each transcript is different, the associated biological functions are conserved in different cancers.

First, we focused on GO terms that are consistently enriched in more than 10 cancers and conservatively associated with the corresponding transcript. We investigated the enriched GO terms associated with ENST00000335181 encoding PKM2. As shown in Figure 1B, glycolytic process, hypoxia response, NADH regeneration pathways are enriched with upregulated genes in patients with high expression of ENST00000335181. This is expected since it reflects the key enzymatic role in glycolysis of PKM2. On the other hand, ATP synthesis, mitochondrial respiratory process, oxidative phosphorylation pathways are enriched with downregulated genes, which probably indicated the shift from oxidative phosphorylation to glycolysis that is well known as the Warburg effect in cancer. Moreover, the upregulated genes were also enriched in pathways associated with the cell morphology such as cell motility, cell migration, cell adhesion and cell junction pathways. These could be linked to the tumorigenesis role of PKM2^7,8,26,27^. Interestingly, several RNA processing related pathways, translational initiation pathway and pathways related to protein localization were also enriched with down-regulated genes in cancer patients with high expression of ENST00000335181. These pathways were rarely associated with the biological function of *PKM* in previous studies and appeared as commonly enriched GO terms in the same analysis for other three transcripts. Hence, studying alternative splicing processes of *PKM* in different cancers provided further understanding about the role of *PKM* in cancer progression.

Second, we investigated the function of other three transcripts of *PKM* whose functions have not been known. We found that many of the enriched GO terms associated with these three transcripts are similar to the GO terms associated with ENST00000335181 (Figure 1B). However, the GO terms associated with the ENST00000561609 followed the same direction with those of ENST00000335181, of which followed the opposite direction with the GO terms associated with both ENST00000389093 and ENST00000568883. In total, 27 different GO terms, e.g. oxidative phosphorylation, translational initiation and RNA catabolic processes, are enriched with genes that are downregulated with both ENST00000335181 and ENST00000561609, and genes that are upregulated with both ENST00000389093 and ENST00000568883.

We also compared the DEGs identified by comparing high and low expression of each transcript and observed similar results based on the directionality and overlap of the DEGs. For instance, in KIRC, we identified 3162 and 6592 DEGs when comparing the high and low expression of ENST00000335181 and ENST00000561609, which both exhibited favorable prognostic indications, respectively. We found that the two sets of DEGs has a significant overlap (*n* = 2010; hypergeometric distribution test, *p* < 1.11e-16) and the concordance score of these overlapped genes (using directionality of the DEGs) is 99.06%. We also identified 6541 and 6885 DEGs when comparing the high and low expression of ENST00000389093 and ENST00000568883 transcript pair, which both exhibited unfavorable prognostic effect in KIRC, respectively. Notably, we found that the overlap between the DEGs of the transcripts is 5469 (hypergeometric distribution test, *p* < 1.11e-16) and the concordance score is 100%. On the other hand, we investigated the overlap between DEGs associated with the transcripts exhibiting the opposite prognostic effect and found that there is no statistically significant overlap. For instance, in KIRC, the concordance score of the overlapped DEGs between transcripts with opposite prognostic indications were between 0.10% and ∼20% (Table S4). Similar scenarios were also observed in CESC, PAAD, BRCA, and COAD (Table S5-8). Our results suggested that both ENST00000335181 and ENST00000561609 have similar biological functions and both have opposite biological functions compared to the other pair of transcripts including ENST00000568883 and ENST00000389093.

### Validation of the prognostic effect in independent KIRC cohort

We performed a survival analysis for these four transcripts in 100 KIRC patients involved in an independent Japanese study^28^. As shown in Figure 2A, the high expression of both ENST00000335181 and ENST00000561609 are significantly (log-rank *p* < 0.05) associated with the favorable survival of patients whereas the high expression of ENST00000389093 and ENST00000568883 are significantly associated with an unfavorable survival of patients. Our analysis indicated that the former pair is favorable prognostic transcripts and the latter pair is unfavorable prognostic transcripts in KIRC and it agrees with our results based on the TCGA KIRC cohort. We also identified the DEGs by comparing the patients with top 25% high expression and bottom 25% low expression of each transcript in the Japanese KIRC cohort. We performed a GO terms enrichment analysis for the DEGs identified by each transcript, and summarized the results in Figure S1. We found that the mitochondrial respiratory process, ATP synthesis, oxidative phosphorylation, ribonucleotide metabolic process, and purine nucleotide metabolic process pathways are enriched with downregulated genes in patients with high expression of ENST00000561609 and upregulated genes in patients with high expression of ENST00000389093 and ENST00000568883. We also observed that the chromatin modification and histone modification pathways are enriched with upregulated genes in patients with high expression of ENST00000561609 and downregulated genes in patients with high expression of ENST00000389093 and ENST00000568883. We observed that the biological functions associated with DEGs identified by comparing high and low expression of each transcript in the Japanese KIRC cohort agree with the associations identified in the TCGA KIRC cohort.

**Figure 2.**
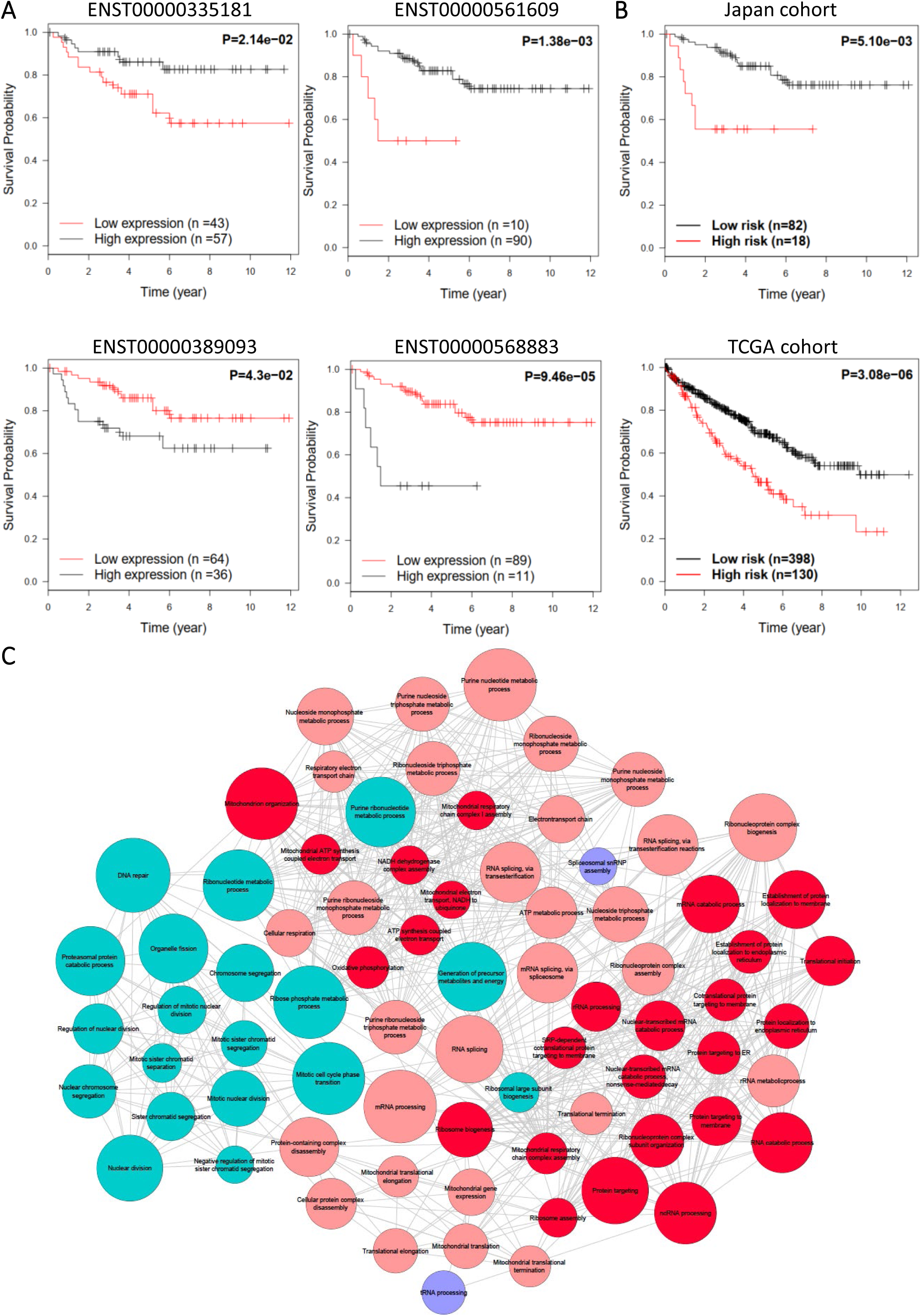
Validation of the prognostic effect and biological function of transcripts in an independent KIRC cohort. (A) The Kaplan Meier plots for the samples classified by the high andlow expressionoftranscripts including ENST00000335181, ENST00000561609, ENST00000389093 and ENST00000568883 in Japanese cohort. (B) The Kaplan Meier plots for the samples classified by the prognostic signature in TCGA and Japanese KIRC cohorts. (C) Network plot of enriched GO terms for DEGs between TCGA and Japanese KIRC cohorts. Sizes of the nodes are correlated to the corresponding total number of genes, and connections between the nodes indicate the significant overlaps (hypergeometric distribution test; FDR < 1.0e-05) between the genes of the corresponding GO terms. Nodes in red, blue and purple indicate GO terms that enriched in both cohorts, only in Japanese KIRC cohort, and only in TCGA KIRC cohort, respectively. The nodes highlighted in dark red indicate the common GO terms associated with all four transcripts reported in this study.

To investigate whether these transcripts regulate the similar genes in two different cohorts, we compared the DEGs by comparing the expression of each transcripts in patients with the top 25% high expression and bottom 25% low expression in Japanese and TCGA KIRC cohorts. For fair comparison, we selected the top 20% of the DEGs (*n* = 2694) in TCGA and Japanese cohorts and checked their overlap between genes. We found that the number of overlapped DEGs identified for ENST00000561609, ENST00000389093 and ENST00000568883 are 1370, 1499 and 1449 (hypergeometric distribution test, *p* < 1.11e-16) and the concordance scores between the cohorts are 100%, 99.93% and 99.86%, respectively (Table S9). Our analysis indicated that the biological functions associated with each of these three transcripts are highly conserved in independent KIRC cohorts. However, we found that the number of overlapping DEGs identified with the transcript ENST00000335181 in both cohorts is relatively small (*n* = 546; hypergeometric distribution test, *p* **≈** 1) and the concordance score is 75.46%, which also indicates the differences between the two cohorts. Such differences may be explained by the dietary and geographical differences between the two independent cohorts.

### Combined prognostic signature for KIRC

Based on the highly conserved prognostic effects of the *PKM* transcripts in two independent KIRC cohorts, there is likely to be different molecular subtypes among KIRC patients with opposite expression patterns of the transcripts highlighted in this study. Thus, we extracted a prognostic signature based on the expression value of these four transcripts (see method). In brief, if more than half of the transcripts indicates an unfavorable prognosis, the patient is classified as high-risk and otherwise as low-risk. Using this rule, we observed significantly different overall survival (log-rank *p* < 0.01) between high- and low-risk groups in both TCGA and Japanese KIRC cohorts as shown in Figure 2B.

To investigate whether these two molecular subtypes identified in both cohorts exhibited similar biological differences, we extracted the top 20% most significant DEGs (*n* = 2694) between high-risk and low-risk groups in the TCGA and Japanese cohorts. The two lists of DEGs had significant overlap (*n* = 1516; hypergeometric distribution test, *p* < 1.11e-16) and the concordance score was 100%. In addition, we identified 57 and 74 GO terms that are significantly enriched with upregulated genes (FDR < 1.0e-05) in the high-risk group of the TCGA and Japanese cohorts, respectively (Figure 2C, Table S10). Interestingly, we found that 55 of these enriched GO terms are common in both cohorts and the molecular subtypes identified by our analysis have consistent biological differences. Moreover, 26 of the 27 GO terms that are significantly associated with the four transcripts (shown in Figure 1B, e.g. oxidative phosphorylation, translational initiation and RNA catabolic process) are also among the overlapped enriched GO terms, which further indicated that the molecular subtypes are functionally related to the four key transcripts of *PKM* identified in this study.

### Discovery of the protein products of the prognostic transcripts

To investigate and compare the protein products of the three novel transcripts including ENST00000561609, ENST00000389093 and ENST00000568883, whose functions were previously unknown compared to the function of ENST00000335181 (encoding PKM2), we first aligned their amino acid (AA) sequences (Table S11). We observed that ENST00000335181 has the longest AA sequence with 531aa, followed by the protein products of ENST00000561609, ENST00000389093, and ENST00000568883, which are 485aa, 457aa, and 366aa, respectively. As shown in Figure 3A, we found that proteins of ENST00000389093 and ENST00000568883 miss a part of the A1 and B domains (59-132aa and 41-205aa) in PKM2, which may affect the formation of dimer^29^. We also found that the protein product of ENST00000561609 is shorter than PKM2, missing amino acid residues 486-531aa from PKM2 which is a part of the C-domain participating in the formation of tetramer^29^. This implicated that the protein encoded by ENST00000561609 may have no tetrameric formation. In addition, there is part of the AA sequence, 389-433aa, of the protein product of ENST00000561609 and ENST00000568883 resembles PKM1 protein rather than PKM2. In this part, K433 is the fructose 1,6-bisphosphate (FBP) binding site in PKM2, which activates the association of monomer to form the tetrameric^30^. However, PKM1 does not bind FBP due to AA difference at the FBP binding pocket and it naturally exists as a stable tetramer that has high constitutive activity^31^. In addition, all protein products of the three transcripts have K270, which is the active site, binding to phosphoenolpyruvate (PEP).

**Figure 3.**
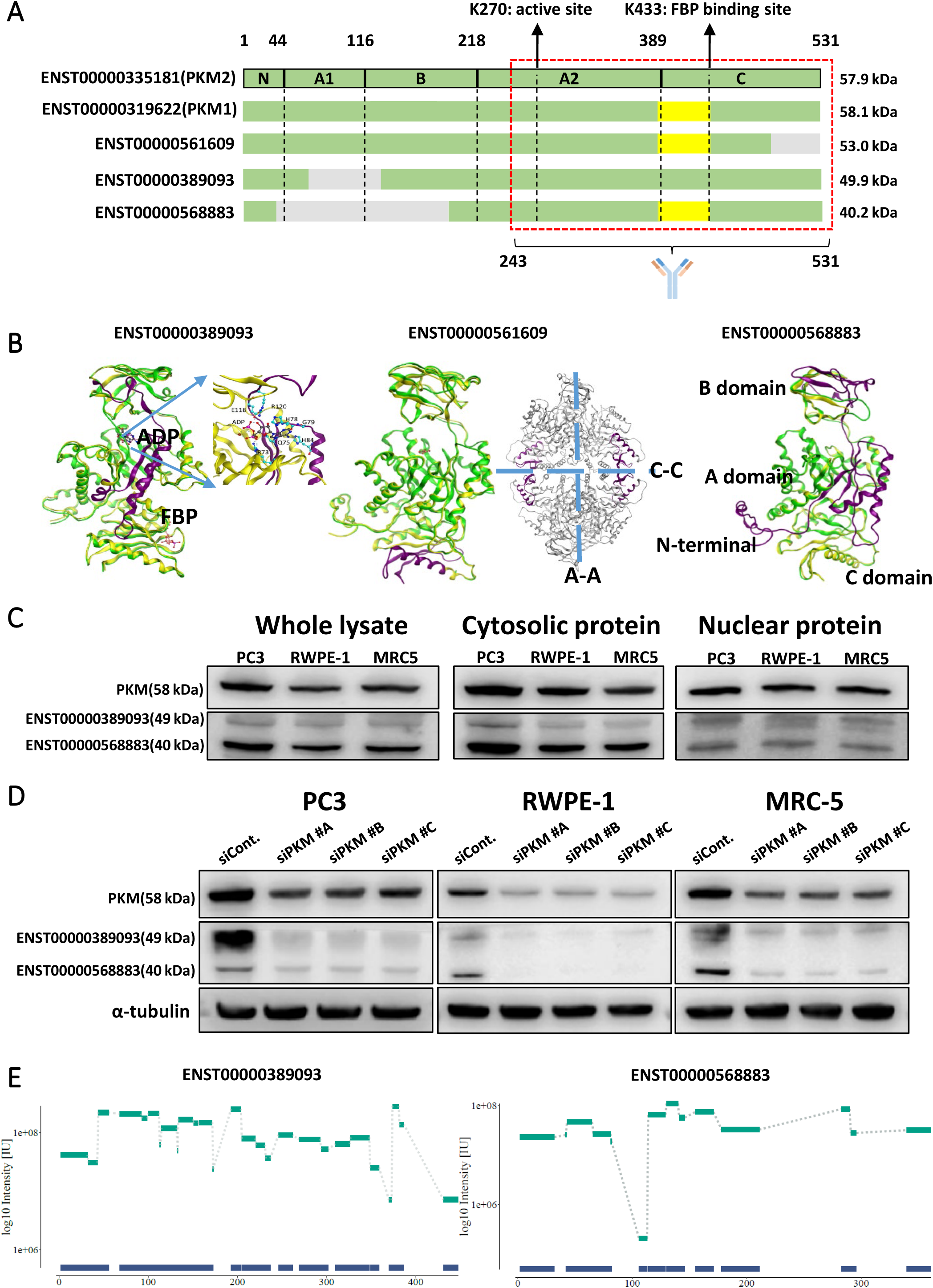
Protein products of the functional alternatively spliced PKM transcripts. (A) Alignment of amino acid sequence of the four functional different transcripts and the transcript encoding PKM1. The gray color denotes missing part compared to PKM2. The yellow color denotes a subsequence that is specific to PKM1. (B) Homology modelling predicted structures of the protein products of ENST00000561609, ENST00000389093 and ENST00000568883. (C) Western blots for the proteins encoded by the transcript ENST00000389093 and ENST00000568883. (D) Western blots showing the protein level of PKM2 and protein products of ENST00000389093 and ENST00000568883 with siRNA and negative control. (E) Peptides detected using LC-MS/MS aligned with amino-acid sequence of respective transcript products on the x-axis and intensity on the y-axis.

Furthermore, we constructed the homology models of ENST00000561609, ENST00000389093 and ENST00000568883 to obtain the protein structure information. When compared to the PKM2 structure, we found that the ENST00000389093 structure is missing the catalytic site for ADP binding (59-132aa) and several AA residues, including R73, Q75, H78, G79, H80, E118, and R120 from the missing part are in close contact with ADP (Figure 3B). Instead, the ENST00000389093 structure forms a newly ordered loop going straight through the ADP binding site, and whether this loop is able to coordinate the ADP binding is still unknown. On the other hand, we observed that the PEP and the FBP binding site in ENST00000389093 structure is fully maintained as in the PKM2 structure, and the tetramer binding interface is kept the same as in PKM2 structure. The protein product of ENST000000561609 shares the exact the same AA sequence as in PKM1, but missing the AAs from 486 to 531 in PKM1. By overlapping the structure of the protein product of ENST000000561609 with PKM1, we found that the protein maintains well defined ADP and PEP binding sites. However, the missing part constitutes part of the C-C binding interface (Figure 3B). Therefore, whether the ENST000000561609 functions as a monomer or active tetramer needs further investigation. Comparing the protein of ENST000000568883 with PKM1, we observed that it is missing large part of both A and B domain as well as the whole N-terminal part. This led to a loss of a large part in the ADP binding site and the binding interface, whereas the PEP binding site in ENST000000568883 structure (Figure 3B) was kept.

As we have shown that the AA sequence of the protein products of ENST00000561609, ENST00000389093 and ENST00000568883 resemble different part of either PKM1 or PKM2, it is difficult to stratify them based on the AA sequence only. However, we observed that all of these transcripts have different length of AA sequences, and different protein masses. The protein masses for PKM1 and PKM2 are 58.1 kDa and 57.9 kDa, respectively, while the mass for protein products of ENST00000561609, ENST00000389093 and ENST00000568883 are 53.0 kDa, 49.9 kDa and 40.2 kDa, respectively. Therefore, we separated these proteins based on their mass differences using sodium dodecyl sulfate (SDS) gel electrophoresis and evaluated them by Western blot. We used an antibody targeting the latter half of PKM2 for the detection of the other three transcripts since they shared a large portion of AA sequence in those areas as shown in Figure 3A. We found that there are different bands that appearing around 49 kDa and 40 kDa in the western blots (Figure 3C) in three different human cell lines, which is in very good agreement with the putative mass of protein products from ENST00000389093 and ENST00000568883, respectively. We also found the two bands in the same location in the western blots of the nuclear proteins, which means these two proteins may also play important roles in cell nucleus as PKM2.

Although we observed the bands that potentially represent the protein products of ENST00000389093 and ENST0000568883, there is still chance that these band are shown because of the non-specific binding of the antibody. To further validate whether the bands we identified are encoded by ENST00000389093 and ENST00000568883, we used three different siRNAs to inhibit the expression all protein isoforms of *PKM*. As shown in Figure 3D, the cellular *PKM* level is decreased with the siRNA transferred to the cells. In addition, we found the bands located at 49 kDa and 40 kDa also significantly decreased. This indicated that the two bands we identified are encoded by *PKM*. Moreover, we manually cut the gel with protein in PC3 from 37 kDa to 50 kDa based on the marker and separated it into three horizontal slices. Subsequently, we subjected the samples to enzymatic digestion and extracted peptides for analysis in mass spectrometry (MS). Consequently, we detected signals of peptides on both the top (49.9 kDa) and bottom (40.2 kDa) slices. As shown in Figure 3E, both of these slices showed high MS intensity with good peptide coverage, proving that the bands we identified are related to the corresponding transcripts. With respect to the one from ENST00000561609, we could not visually separate the bands for PKM1 and PKM2 since they have a similar mass with PKM1 and PKM2.

### The expression of prognostic transcripts in normal tissues and cancers

To further investigate the prevalence of the discovered proteins in different cancers, we summarized the mRNA expression of the four transcripts as well as the transcript for PKM1 in all cancers. As shown in Figure S2, the three newly discovered prognostic transcripts of *PKM*, including ENST00000561609, ENST00000389093 and ENST0000568883 have higher expressions compared to the transcript of PKM1 in the TCGA dataset. In addition, all these three transcripts showed significant inter-cancer variations, while the transcript encoding PKM2, had a house-keeping expression profile in all cancers.

We also investigated the mRNA expression of the PKM transcripts in the corresponding normal tissues. In this context, we presented the mRNA expression of these transcripts in the matched normal tissue in TCGA dataset (Figure S3), as well as in GTEx database (Figure S4). We found clear tissue specific pattern based on the expression of ENST00000561609 and ENST0000568883. We also found that the expressions of ENST00000389093 are low in all normal tissues compared to other transcripts, but still in the same order of magnitude compared to the expression of PKM1. Therefore, we concluded that the three prognostic transcripts discovered in this study are expressed in different normal and cancer tissues, and their expression is significantly increased in different cancers.

## DISCUSSION

Several studies have been performed for studying the functional role of *PKM* in cancer metabolism, mainly focusing on PKM1 and PKM2 isoforms. With the development of bioinformatics tools for the analysis of the next-generation sequencing data, such as RSEM^32^ and Kallisto^33^, now it is possible to perform systematic studies for revealing the functional role of transcripts in cancer progression. In this study, we performed a transcript level analysis of *PKM* and found that four of them, including PKM2 but not for PKM1, could play a key role in KIRC progression. We found that mRNA expression of these four transcripts is also significantly associated with the survival of the patients with different cancers. Next, we investigated the functional role of each transcript, identified the associated biological functions and validated their prognostic effect in an independent KIRC cohort. We also identified a signature to stratify patients with kidney cancer into two groups with distinct biological features and survival. Finally, we characterize for the first time the protein products of these key transcripts using Western blots and mass spectrometry-based proteomics data and showed the relevance of these transcripts in different normal tissues and cancer.

Previous studies reported that the ratio between PKM1 and PKM2 isoforms plays a key role in cancer progression^34-38^. In our study, we found PKM1 is not strongly associated with the survival of cancer patients as it has been reported^18^. Based on our analysis, we observed that the disagreement between the studies may be explained by the differences in transcriptomic quantification methods used in the analysis of the data. For instance, a recent study quantified the mRNA level of PKM1 and PKM2 using RT-PCR and reported their association in cancer^39^, but the primer they have used for RT-PCR could also bind to ENST00000568883 and ENST00000561609. Thus, the RNA level suggested for PKM1 transcript may actually be the sum of all three transcripts rather than just PKM1. In addition, we also found that the RNA level of PKM1 (<5%) is very low compared to PKM2 (∼95%) which is in good agreement with proteomics data reported earlier^40^, and it has the same magnitude as ENST00000568883 and ENST00000561609 (Table S1). In this context, it is very likely that ENST00000568883 or ENST00000561609 which showed prognostic effect in our study may also play a key role in tumor metabolism.

We also showed the protein products of these two transcripts using Western blots and validated their presence by MS after the identification of functional alternatively spliced *PKM* transcripts. It is quite difficult to distinguish these transcripts and PKM2 since they shared the majority of their nucleotide and AA sequences. For instance, the antibody we used in this study is designed to specifically target the PKM2 protein, but it also bound the protein products of ENST00000568883 and ENST00000389093. Therefore, this might be a potential explanation for the contradicting prognostic effect related to PKM2 in different studies, and there may be a need to revisit some of the previous studies to investigate all isoforms of *PKM*.

In conclusion, we revealed the functional role of the three alternatively spliced *PKM* transcripts in KIRC and different cancers based on an integrative systems analysis. Our study may be considered as a primer for the future studies focusing on the biological and oncological role of alternatively spliced gene transcripts of not only *PKM* but also other gene targets. Such analysis may allow for the discovery of the right protein product which could be effectively targeted using pharmaceutical agents.

## Acknowledgements

The study is funded by The Knut and Alice Wallenberg Foundation.

## Materials and Methods

### Data and preprocessing

The TCGA transcript-expression level profiles (TPM and count values) of 25 cancer-types with more than 100 patients and excluding LGG for the same reason as in our previous study^3^ were download from https://osf.io/gqrz9^41^ on November 27, 2018, which were quantified by Kallisto^33^ based on the GENCODE reference transcriptome (version 24) (Ensembl 83 (GRCh38.P5)). The clinical information of TCGA samples was downloaded through R package TCGAbiolinks^42^. The whole-exome sequences data of 100 KIRC samples of patients from Japanese cohort^28^ were download from European Genome-phenome Archive (accession number: EGAS00001000509). BEDTools^43^ was used for converting BAM to FASTQ file. Kallisto was used for estimating the count and TPM values of transcripts based on the same reference transcriptome of TCGA data. The sum value of the multiple transcripts of a gene was used as the expression value of this gene. The genes with average TPM values >1 across patients in each cancer were analyzed. The transcript-expression level data of GTEx with 31 normal human tissues was downloaded from https://xenabrowser.net/datapages/?hub=https://toil.xenahubs.net:443^44^.

### Survival analysis

Based on the TPM value of each transcript or gene, we classified the patients into two groups and examined their prognoses. Survival curves were estimated by the Kaplan-Meier method and compared by log-rank test. To choose the best TPM cutoffs for grouping the patients most significantly, all TPM values from the 10^th^ to 90^th^ percentiles used to group the patients, significant differences in the survival outcomes of the two groups were examined and the value yielding the lowest log-rank *p* value was selected.

For retrieving prognostic signature, we used the expression cutoff obtained in the individual survival analysis for each of the four transcripts which could classify the patients into two groups with significantly different prognoses. In TCGA cohort, if the expression of ENST00000335181 or ENST00000561609 is less than 476.35 or 0.69 in a sample, this sample would be classified into high-risk group, otherwise, low-risk group. On the other hand, if the expression value of ENST00000389093 or ENST00000568883 is higher than 18.18 or 13.74 in a sample, this sample will be classified into high-risk group, otherwise, low-risk group. Similarly, the cutoffs of the four transcripts are 815.84, 0.33, 7.90 and 6.63 in Japanese cohort. Therefore, we classified the samples of the two different cohorts into the high-risk group when at least two transcripts are higher or lower than the corresponding cutoffs.

### Differential expression analysis

DESeq2^45^ was used to identify DEGs between two groups. The raw count values of genes were used as input of DESeq2. The Benjamini-Hochberg procedure was used to estimate FDR.

### Overlapping of two lists of DEGs

If DEG list 1 with *L*_*1*_ genes and DEG list 2 with *L*_*2*_ genes have *k* overlapping genes, among which *s* genes shows the same directions (up or down-regulation) in the two DEGs lists, the probability of observing at least *s* consistent genes by chance can be calculated according to the following cumulative hypergeometric distribution model:

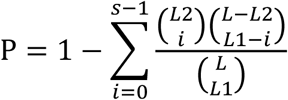

where *L* represents the number of the background genes commonly detected in the datasets from which the DEGs are extracted. The two DEG lists were considered to be significantly overlapping if P<0.05.

The concordance score *s*/*k* is used to evaluate the consistency of DEGs between the two lists. Obviously, the score ranges from 0 to 1, and the higher concordance score suggests the better consistency of two lists of DEGs.

### Functional enrichment analysis

GO enrichment was performed by the enrichGo function in R package ClusterProfiler^46^, in which the hypergeometric distribution was used to calculate the statistical significance of biological pathways enriched with DEGs of interest.

### Hierarchical Clustering

Log-rank *p* values were hierarchically clustered by Spearman correlation distance and Ward linkage method (ward.D2). Negative log 10 transformation of *p* values was performed before clustering.

### Western blots

Cell lysate was extracted with CelLytic M (C2978, Sigma-Aldrich) lysis buffer. Cytosolic and nucleus protein was extracted with Nuclear Extraction Kit (ab113474, abcam). Isolation and protein extraction of mitochondria was performed by Mitochondria Isolation Kit for Cultured Cells (ab110170. abcam). Proteins were separated by Mini-PROTEAN® TGX™ Precast Gels (Bio-Rad, CA, USA) and transferred using Trans-Blot(r) Turbo™ Transfer System (Bio-Rad, CA, USA). PKM2 antibody (ab137791, abcam) was used for primary antibody overnight. Two PKM isotype band (49.9 kDa and 40.2 kDa) were detected with ImageQuanat^tm^ LAS 500 (29-0050-63, GE) for 5 min exposure.

### Cell culture and siRNA transfection

All cells were cultured followed by ATCC instruction. PC3 cells culture media formulation is F12K Nutrient mix supplemented with 10% FBS and 1% Penicillin/Streptomycin, MRC5 cells cultured with DMEM with 10% FBS and 1% Penicillin/Streptomycin, and RWPE-1 cells was cultured with Keratinocyte Serum Free Medium (K-SFM) supplemented with Bovine Pituitary Extract (BPE) and human recombinant Epidermal Growth Factor (EGF) (Kit Catalog Number 17005-042). For siRNA treatment, 400,000cells were seeded to 6 well plate. After 24 hr of cell seeding 25pmol siRNA was transfected by Lipofectamine(®) RNAiMAX (13778-075 Invitrogen) for two days.

### Sample preparation for mass spectrometry analysis

The gel pieces were subjected to in-gel digestion as descried by Shevchenko, et al. ^47^, with some adjustments. Reduction was performed by addition of 10 mM dithiothreitol and incubation at 56 °C for 30 min. The samples were alkylated by addition of 55 mM 2-chloroacetamide and incubation shielded from light for 20 min at room temperature. Tryptic digestion was performed overnight at 37 °C after addition of trypsin solution containing 13 ng/µl proteomics grade porcine trypsin (Sigma Aldrich, St Louis, MO, USA), 100 mM ammonium bicarbonate, 10% acetonitrile (ACN). The peptides were then extracted by addition of 100 µl extraction buffer to each sample (1:2, 5% fomic acid (FA)/ACN). The extracted peptides were transferred to HPLC-vials and dried using vacuum centrifugation. The peptides were then resuspended in 60 µl 3% ACN, 0.1% FA and analyzed by liquid chromatography (LC)-MS/MS.

### LC-MS/MS analysis

PC3 cell lysate was prepared with CelLytic M (C2978, Sigma-Aldrich) lysis buffer. SDS PAGE separated 60µg of PC3 cell lysate per well with Precision Plus Protein Standards ladder (1610374, Biorad). Gel pieces were cut by razor blade, three pieces between 37 and 50 kDa ladder indicated.

The samples were analyzed using a Thermo Scientific Q Exactive HF (Thermo Fisher Scientific, Waltham, MA, USA) online connected to a Dionex Ultimate 3000 UHPLC-system (Thermo Fisher Scientific) equipped with a reverse phase trap column (Acclaim PepMap 100, 75 µm x 2 cm, 3 µm, 100 Å; Thermo Fisher Scientific) and 50 cm analytical column (EASY-Spray, 75 µm x 50 cm, 2 µm, 100 Å; Thermo Fisher Scientific). 10 µl of each sample was injected for analysis and the peptides were separated over an 85 min run using a 60 min linear LC-gradient and sprayed directly into the mass spectrometer using the EASY-Spray ion source. The solvents used for the LC-gradient were 3% ACN, 0.1% FA (solvent A) and 95% ACN, 0.1% FA (solvent B). The flow rate of the system was set to 300 nl/min and the gradient used was as follows: 5% solvent B for 3 min, 5-35% solvent B within 60 min, 35-90% solvent B within 5 min, 90% solvent B for 7 min, 90-5% solvent B within 0.1 min, 5% solvent B for 10 min. The mass spectrometer was set to operate using a Top10 MS method with a full scan resolution of 60,000 (mass range: 400-1,200 m/z, AGC: 3e6) and a MS/MS resolution of 30,000 (AGC: 1e5). The normalized collision energy was set to 30.

### Data analysis of LC-MS/MS results

The raw files obtained from the LC-MS/MS experiment were analyzed using MaxQuant (version 1.6.1.0)^48^ implementing Andromeda^49^ to search the MS/MS data against the Ensembl Homo sapiens database (version 83.38, all protein coding transcripts from the primary assembly) as well as a separate database with the two distinct target sequences (ENST00000389093 and ENST00000568883) a list of common contaminants. Trypsin/P was used for cleavage specificity with up to two missed cleavages. Oxidation (M) was used as a variable modification while carbamidomethylation (C) was used as a fixed modification. The peptide and protein FDR were set to 1% and the minimum peptide length was set to seven amino acids. The presence of the target proteins was assessed by evaluating the identification of unique peptides specific to the proteins in the different samples.

### Homology model

The homology models were built using StructurePrediction panel in Schrödinger Suite (Schrödinger, LLC, New York, NY). The ClustralW method was used to align the target and template sequences in Prime, the energy-based was selected for model building method, and homo-multimer was selected for multi-template model type. The homology model of ENST00000561609 was built based on the PKM2 crystal structure (PDB ID: 5X1W), as ENST00000561609 shares 96% sequence similarity to PKM2, compare to 91% to PKM1. ENST00000389093 and ENST00000568883 share higher sequence similarity to PKM1, with 100% and 92% correspondingly. These two homology models were built based on the PKM1 crystal structure (PDB ID: 3SRF).

## Supplementary Tables

Table S1. The average expression of PKM and its 14 transcripts in 25 cancer-types

Table S2. Log-rank *p* value of PKM and its 14 transcripts in 25 cancer-types

Table S3. The enriched of GO terms with DEGs for all transcripts in 25 cancer-types

Table S4. Overlapping of DEGs between transcripts in KIRC

Table S5. Overlapping of DEGs between transcripts in CESC

Table S6. Overlapping of DEGs between transcripts in PAAD

Table S7. Overlapping of DEGs between transcripts in BRCA

Table S8. Overlapping of DEGs between transcripts in COAD

Table S9. Overlapping of DEGs between TCGA and Japanese cohorts for four transcripts

Table S10. The enriched GO terms for the DEGs identified from TCGA and Japanese cohorts

Table S11. The alignment of amino acid sequences encoded by ENST00000319622, ENST00000335181,ENST00000561609,ENST00000389093and ENST00000568883

**Figure S1.**
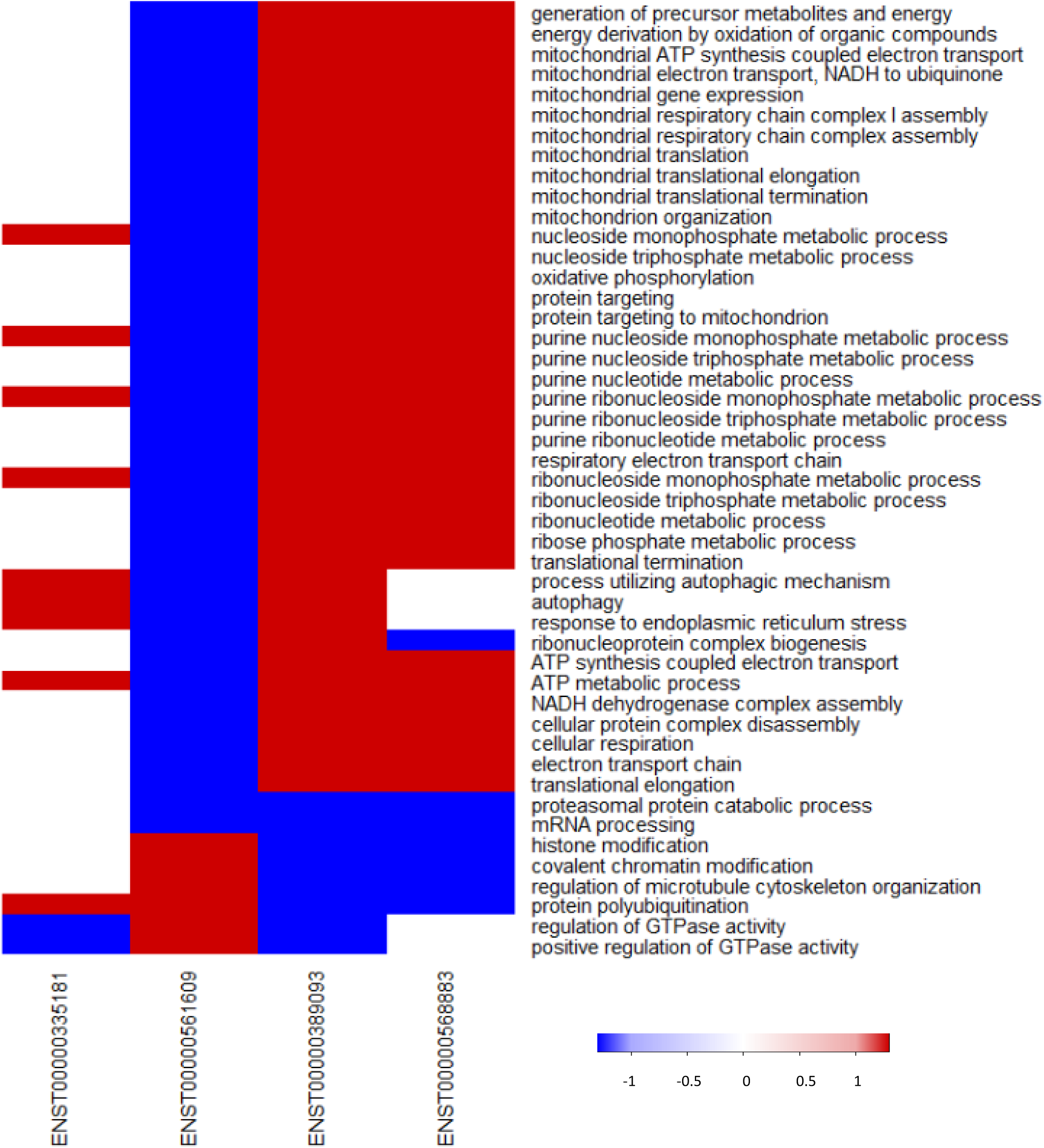
Heat map for the enriched GO terms for four functional transcripts in Japanese KIRC cohort.

**Figure S2.**
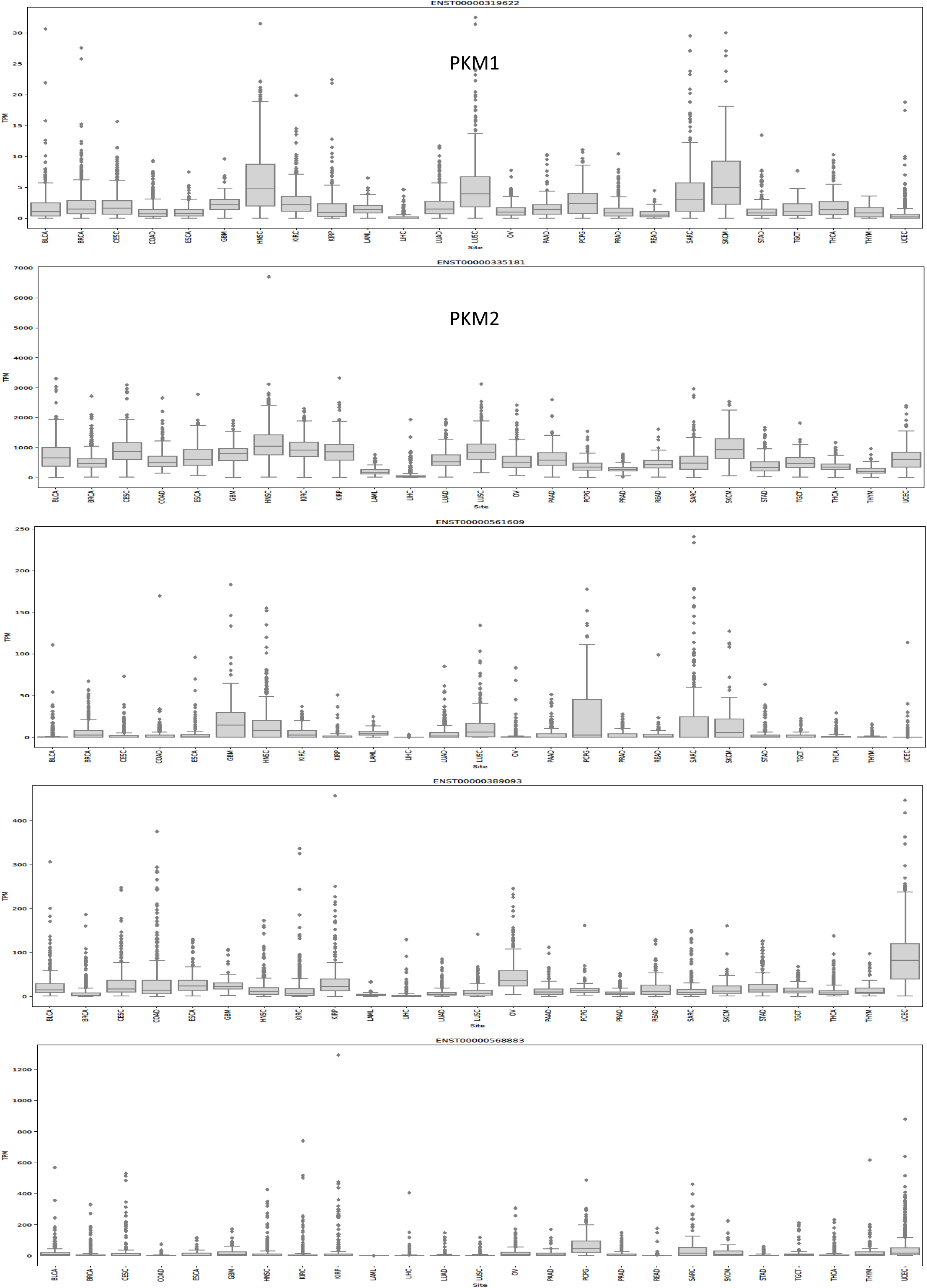
Boxplots showing the RNA expression of transcripts in TCGA tumor samples.

**Figure S3.**
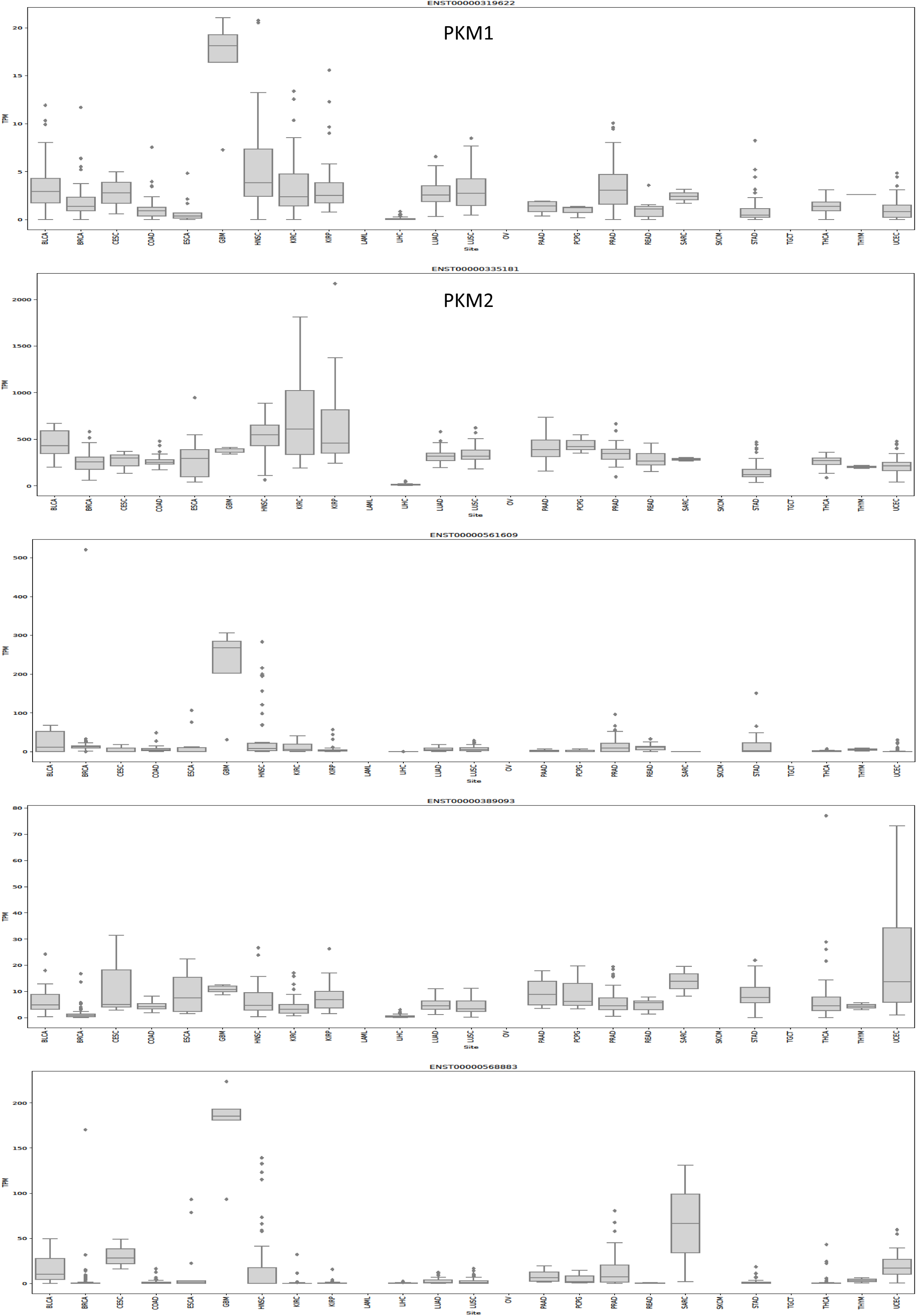
Boxplots showing the RNA expression of transcripts in TCGA normal samples.

**Figure S4.**
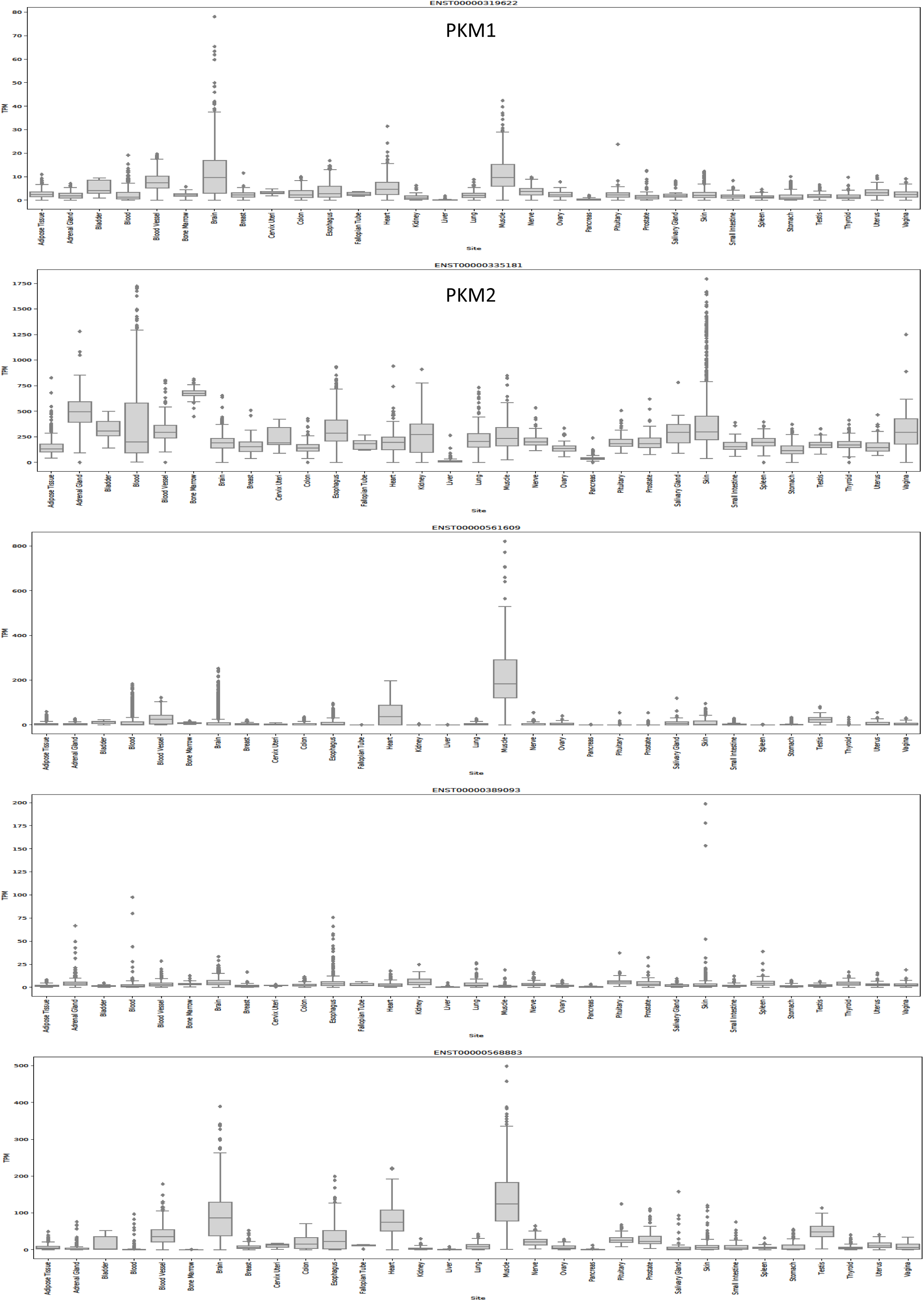
Boxplots showing the RNA expression of transcripts in GTEx normal samples.

## References

1 Allen, A. E. & Locasale, J. W. Glucose Metabolism in Cancer: The Saga of Pyruvate Kinase Continues. Cancer Cell 33, 337–339, doi:10.1016/j.ccell.2018.02.008 (2018).

2 Dong, G. et al. PKM2 and cancer: The function of PKM2 beyond glycolysis. Oncol Lett 11, 1980–1986, doi:10.3892/ol.2016.4168 (2016).

3 Uhlen, M. et al. A pathology atlas of the human cancer transcriptome. Science 357, doi:10.1126/science.aan2507 (2017).

4 Dayton, T. L., Jacks, T. & Vander Heiden, M. G. PKM2, cancer metabolism, and the road ahead. EMBO Rep 17, 1721–1730, doi:10.15252/embr.201643300 (2016).

5 Morita, M. et al. PKM1 Confers Metabolic Advantages and Promotes Cell-Autonomous Tumor Cell Growth. Cancer Cell 33, 355–367 e357, doi:10.1016/j.ccell.2018.02.004 (2018).

6 Chiavarina, B. et al. Pyruvate kinase expression (PKM1 and PKM2) in cancer-associated fibroblasts drives stromal nutrient production and tumor growth. Cancer Biol Ther 12, 1101–1113, doi:10.4161/cbt.12.12.18703 (2011).

7 Jiang, Y. et al. PKM2 regulates chromosome segregation and mitosis progression of tumor cells. Mol Cell 53, 75–87, doi:10.1016/j.molcel.2013.11.001 (2014).

8 Yang, W. et al. PKM2 phosphorylates histone H3 and promotes gene transcription and tumorigenesis. Cell 150, 685–696, doi:10.1016/j.cell.2012.07.018 (2012).

9 Gao, X., Wang, H., Yang, J. J., Liu, X. & Liu, Z. R. Pyruvate kinase M2 regulates gene transcription by acting as a protein kinase. Mol Cell 45, 598–609, doi:10.1016/j.molcel.2012.01.001 (2012).

10 Yang, W. et al. Nuclear PKM2 regulates beta-catenin transactivation upon EGFR activation. Nature 480, 118–122, doi:10.1038/nature10598 (2011).

11 Gupta, A. et al. PAK2-c-Myc-PKM2 axis plays an essential role in head and neck oncogenesis via regulating Warburg effect. Cell Death Dis 9, 825, doi:10.1038/s41419-018-0887-0 (2018).

12 Cheng, T. Y. et al. Pyruvate kinase M2 promotes pancreatic ductal adenocarcinoma invasion and metastasis through phosphorylation and stabilization of PAK2 protein. Oncogene 37, 1730–1742, doi:10.1038/s41388-017-0086-y (2018).

13 Yu, G. et al. PKM2 regulates neural invasion of and predicts poor prognosis for human hilar cholangiocarcinoma. Mol Cancer 14, 193, doi:10.1186/s12943-015-0462-6 (2015).

14 Najera, L. et al. Prognostic implications of markers of the metabolic phenotype in human cutaneous melanoma. Br J Dermatol, doi:10.1111/bjd.17513 (2018).

15 Lv, W. W. et al. Effects of PKM2 on global metabolic changes and prognosis in hepatocellular carcinoma: from gene expression to drug discovery. BMC Cancer 18, 1150, doi:10.1186/s12885-018-5023-0 (2018).

16 Yang, P., Li, Z., Fu, R., Wu, H. & Li, Z. Pyruvate kinase M2 facilitates colon cancer cell migration via the modulation of STAT3 signalling. Cell Signal 26, 1853–1862, doi:10.1016/j.cellsig.2014.03.020 (2014).

17 Wang, Y. et al. Overexpression of pyruvate kinase M2 associates with aggressive clinicopathological features and unfavorable prognosis in oral squamous cell carcinoma. Cancer Biol Ther 16, 839–845, doi:10.1080/15384047.2015.1030551 (2015).

18 Christofk, H. R. et al. The M2 splice isoform of pyruvate kinase is important for cancer metabolism and tumour growth. Nature 452, 230–233, doi:10.1038/nature06734 (2008).

19 Lunt, S. Y. et al. Pyruvate kinase isoform expression alters nucleotide synthesis to impact cell proliferation. Mol Cell 57, 95–107, doi:10.1016/j.molcel.2014.10.027 (2015).

20 Liu, F. et al. PKM2 methylation by CARM1 activates aerobic glycolysis to promote tumorigenesis. Nat Cell Biol 19, 1358–1370, doi:10.1038/ncb3630 (2017).

21 Dayton, T. L. et al. Germline loss of PKM2 promotes metabolic distress and hepatocellular carcinoma. Genes Dev 30, 1020–1033, doi:10.1101/gad.278549.116 (2016).

22 Israelsen, W. J. et al. PKM2 isoform-specific deletion reveals a differential requirement for pyruvate kinase in tumor cells. Cell 155, 397–409, doi:10.1016/j.cell.2013.09.025 (2013).

23 Tech, K. et al. Pyruvate Kinase Inhibits Proliferation during Postnatal Cerebellar Neurogenesis and Suppresses Medulloblastoma Formation. Cancer Res 77, 3217–3230, doi:10.1158/0008-5472.CAN-16-3304 (2017).

24 Wei, L. et al. Oroxylin A activates PKM1/HNF4 alpha to induce hepatoma differentiation and block cancer progression. Cell Death Dis 8, e2944, doi:10.1038/cddis.2017.335 (2017).

25 Taniguchi, K. et al. MicroRNA-124 inhibits cancer cell growth through PTB1/PKM1/PKM2 feedback cascade in colorectal cancer. Cancer Lett 363, 17–27, doi:10.1016/j.canlet.2015.03.026 (2015).

26 Cortes-Cros, M. et al. M2 isoform of pyruvate kinase is dispensable for tumor maintenance and growth. Proc Natl Acad Sci U S A 110, 489–494, doi:10.1073/pnas.1212780110 (2013).

27 Yang, W. et al. ERK1/2-dependent phosphorylation and nuclear translocation of PKM2 promotes the Warburg effect. Nat Cell Biol 14, 1295–1304, doi:10.1038/ncb2629 (2012).

28 Sato, Y. et al. Integrated molecular analysis of clear-cell renal cell carcinoma. Nat Genet 45, 860–867, doi:10.1038/ng.2699 (2013).

29 Wu, S. & Le, H. Dual roles of PKM2 in cancer metabolism. Acta Biochim Biophys Sin (Shanghai) 45, 27–35, doi:10.1093/abbs/gms106 (2013).

30 Ashizawa, K., Willingham, M. C., Liang, C. M. & Cheng, S. Y. In vivo regulation of monomer-tetramer conversion of pyruvate kinase subtype M2 by glucose is mediated via fructose 1,6-bisphosphate. J Biol Chem 266, 16842–16846 (1991).

31 Israelsen, W. J. & Vander Heiden, M. G. Pyruvate kinase: Function, regulation and role in cancer. Semin Cell Dev Biol 43, 43–51, doi:10.1016/j.semcdb.2015.08.004 (2015).

32 Li, B. & Dewey, C. N. RSEM: accurate transcript quantification from RNA-Seq data with or without a reference genome. BMC Bioinformatics 12, 323, doi:10.1186/1471-2105-12-323 (2011).

33 Bray, N. L., Pimentel, H., Melsted, P. & Pachter, L. Near-optimal probabilistic RNA-seq quantification. Nat Biotechnol 34, 525–527, doi:10.1038/nbt.3519 (2016).

34 Mendez-Lucas, A. et al. Glucose Catabolism in Liver Tumors Induced by c-MYC Can Be Sustained by Various PKM1/PKM2 Ratios and Pyruvate Kinase Activities. Cancer Res 77, 4355–4364, doi:10.1158/0008-5472.CAN-17-0498 (2017).

35 Kuranaga, Y. et al. SRSF3, a Splicer of the PKM Gene, Regulates Cell Growth and Maintenance of Cancer-Specific Energy Metabolism in Colon Cancer Cells. Int J Mol Sci 19, doi:10.3390/ijms19103012 (2018).

36 Okazaki, M. et al. The effect of HIF-1alpha and PKM1 expression on acquisition of chemoresistance. Cancer Manag Res 10, 1865–1874, doi:10.2147/CMAR.S166136 (2018).

37 David, C. J., Chen, M., Assanah, M., Canoll, P. & Manley, J. L. HnRNP proteins controlled by c-Myc deregulate pyruvate kinase mRNA splicing in cancer. Nature 463, 364–368, doi:10.1038/nature08697 (2010).

38 Clower, C. V. et al. The alternative splicing repressors hnRNP A1/A2 and PTB influence pyruvate kinase isoform expression and cell metabolism. Proc Natl Acad Sci U S A 107, 1894–1899, doi:10.1073/pnas.0914845107 (2010).

39 Shiroki, T. et al. Enhanced expression of the M2 isoform of pyruvate kinase is involved in gastric cancer development by regulating cancer-specific metabolism. Cancer Sci 108, 931–940, doi:10.1111/cas.13211 (2017).

40 Bluemlein, K. et al. No evidence for a shift in pyruvate kinase PKM1 to PKM2 expression during tumorigenesis. Oncotarget 2, 393–400, doi:10.18632/oncotarget.278 (2011).

41 Tatlow, P. J. & Piccolo, S. R. A cloud-based workflow to quantify transcript-expression levels in public cancer compendia. Sci Rep 6, 39259, doi:10.1038/srep39259 (2016).

42 Colaprico, A. et al. TCGAbiolinks: an R/Bioconductor package for integrative analysis of TCGA data. Nucleic Acids Res 44, e71, doi:10.1093/nar/gkv1507 (2016).

43 Quinlan, A. R. & Hall, I. M. BEDTools: a flexible suite of utilities for comparing genomic features. Bioinformatics 26, 841–842, doi:10.1093/bioinformatics/btq033 (2010).

44 Vivian, J. et al. Toil enables reproducible, open source, big biomedical data analyses. Nat Biotechnol 35, 314–316, doi:10.1038/nbt.3772 (2017).

45 Love, M. I., Huber, W. & Anders, S. Moderated estimation of fold change and dispersion for RNA-seq data with DESeq2. Genome Biol 15, 550, doi:10.1186/s13059-014-0550-8 (2014).

46 Yu, G., Wang, L. G., Han, Y. & He, Q. Y. clusterProfiler: an R package for comparing biological themes among gene clusters. OMICS 16, 284–287, doi:10.1089/omi.2011.0118 (2012).

47 Shevchenko, A., Tomas, H., Havlis, J., Olsen, J. V. & Mann, M. In-gel digestion for mass spectrometric characterization of proteins and proteomes. Nat Protoc 1, 2856-2860, doi:10.1038/nprot.2006.468 (2006).

48 Cox, J. & Mann, M. MaxQuant enables high peptide identification rates, individualized p.p.b.-range mass accuracies and proteome-wide protein quantification. Nat Biotechnol 26, 1367–1372, doi:10.1038/nbt.1511 (2008).

49 Cox, J. et al. Andromeda: a peptide search engine integrated into the MaxQuant environment. J Proteome Res 10, 1794–1805, doi:10.1021/pr101065j (2011).

